# Time dependent genetic analysis links field and controlled environment phenotypes in the model C_4_ grass *Setaria*

**DOI:** 10.1101/083865

**Authors:** Max J. Feldman, Rachel E. Paul, Darshi Banan, Jennifer F. Barrett, Jose Sebastian, Muh-Ching Yee, Hui Jiang, Alexander E. Lipka, Thomas P. Brutnell, José R. Dinneny, Andrew D.B. Leakey, Ivan Baxter

## Abstract

Vertical growth of plants is a dynamic process that is influenced by genetic and environmental factors and has a pronounced effect on overall plant architecture and biomass composition. We have performed twelve controlled growth trials of an interspecific *Setaria italica* x *Setaria viridis* recombinant inbred line population to assess how the genetic architecture of plant height is influenced by developmental queues, water availability and planting density. The nondestructive nature of plant height measurements has enabled us to monitor vertical growth throughout the plant life cycle in both field and controlled environments. We find that plant height is reduced under water limitation and high density planting and affected by growth environment (field vs. growth chamber). The results support a model where plant height is a heritable, polygenic trait and that the major genetic loci that influence plant height function independent of growth environment. The identity and contribution of loci that influence height changes dynamically throughout development and the reduction of growth observed in water limited environments is a consequence of delayed progression through the genetic program which establishes plant height in *Setaria.* In this population, alleles inherited from the weedy *S. viridis* parent act to increase plant height early, whereas a larger number of small effect alleles inherited from the domesticated *S. italica* parent collectively act to increase plant height later in development.

## Introduction

Height is one of the most influential components of plant architecture. Due to ease of measurement, high heritability and agronomic importance, plant height has been an attractive topic for scientific inquiry for over a century (1). Plant investment in vertical growth has adaptive value because it helps a plant to out2compete neighbors for access to solar radiation. However, vertical growth requires a concomitant energetic allocation for construction of stem tissue at the expense of allocation to lateral growth and other processes. Greater height growth also amplifies hydraulic cost (2), and increases the risk of lodging (3). The benefit to cost ratio of such adaptation is likely directly influenced by growth environment (4). Adaptations introgressed into wheat and rice varieties during the Green Revolution that led to reduced height, increased yield, harvest uniformity, improved carbon partitioning and nutrient and water use efficiency. In biomass crops, height has been shown to be positvely correlated with above ground biomass (5,6). Thus, depending on the crop and breeding objective either increased or decreased height may be targeted.

Plant height in the Poaceae, a family of grasses that includes the cereal crops and bioenergy grasses, is a function of internode length and number, the increase of which is terminated at reproductive maturity. Plant height within the grasses is a highly heritable, polygenic trait (7–14). Genetic studies of height in maize, sorghum, sugarcane, wheat, barley and rice have identified well over 100 QTL (7–14) and enabled the cloning of multiple loci that contribute large effects (9,15). Forward genetic screens of mutant populations has also been an effective approach to identify causative genes associated with height (16,17). Although plant height exhibits a strong degree of genetic determinism, it is influenced by environmental factors such as water availability and planting density and exhibits dynamic behavior throughout the plant life cycle (10,18–22).

The strong influence of environment on plant height suggests that multiple signals impinge on the regulation of plant height throughout development. Surprisingly, relatively few studies have assessed how the temporal genetic architecture of this dynamic trait changes through developmental time (14,23). A major challenge of performing such experiments is the difficulties associated with growing large populations in conditions with contrasting environmental variables while capturing high precision measurements of phenotypes at appropriate intervals.

Recent use of modern high2throughput phenotyping (phenomics) technology is beginning to alleviate these technical obstacles (24,25). Utilizing genetic model systems that possess desirable growth attributes and tractable genetics (26–28) in combination with phenomics technology improves our ability to obtain a more holistic systems view of growth through developmental time and in the face of environmental challenge. As a model system, plants in the genus Setaria, possesses many favorable experimental and life history attributes that make temporal genetic analysis of complex traits possible and the genetic dissection of these traits more feasible than in larger evolutionarily related crop plants like maize and sorghum (26,27). In this study we conduct a large scale and multi-environment analysis of plant height. We identify thirty seven QTL associated with plant height throughout development within a *Setaria* recombinant inbred population (RIL) population (29–31) across twelve different experiments varying water availability and planting density. The results of this study are discussed in light of development and environmental variation and suggest a strategy for the fine scale manipulation of plant height in the Poaceae.

## Materials and Methods

### Plant Material

A *Setaria* F7 RIL population comprised of 217 individuals was used for genetic mapping. The RIL population was generated through an interspecific cross between the wild-type *S. viridis* accession, A10, and the domesticated *S. italica* accession, B100 (29–31). Plant height measurements of individuals within this population were collected in six different trials. Four of the six trials were conducted at field sites located at University of Illinois-Champaign. Two field trials designed to assess plant phenotypic response to planting density and water availability were each conducted in both 2013 and 2014. The controlled environment trials were performed in a growth chamber at Carnegie Institute for Science, Stanford, CA and using the Bellweather Phenotyping Facility at Donald Danforth Plant Science Center (32).

### Illinois

Field experiments were conducted on the South Farms at the University of Illinois Urbana-Champaign in summers 2013 and 2014. The field site is rain-fed, tile-drained, has a deep and organically rich Flanagan/Drummer series type soil (33).

For each field experiment, seeds were first germinated in a greenhouse in plug trays (128sq, T.O. Plastics) containing a media mix composed of sphagnum, vermiculite, fine bark, bark ash, gypsum, slow2release nitrogen, dolomitic limestone, and a wetting agent (Met^Mix 360, Sun Gro Horticulture). Plugs were provided with ample water daily by clear water or by fertigation every other day (EXCEL CAL MAG 152525). Seedlings were transplanted by hand into a mechanically tilled field.

### Carnegie Institute for Science

Plants (F7 RIL population) were grown in deep pots (Stuewe & Sons, Oregon, D25L) with one plant per pot (one plant per RIL was phenotyped). Seeds were directly germinated in 14 inch Deepots filled with a soil mixture containing 75% Pro2Mix (PRO-MIX(r) PGX soil, Premier Tech, Canada) and 25% river sand, imbibed in water to pot capacity. Pot capacity, defined as the amount of water that soil in a pot can hold against the pull of gravity, was estimated to be 230 ml. Plants were grown in a growth chamber (12 hours light at 31°C and 12 hours dark at 23°C with constant relative humidity at 54255%). Wel^watered plants were watered back to field capacity once every third day whereas for water deficit experiments, seeds were sown in soil imbibed to pot capacity and no further water was added. To prevent water loss, the bottom of each pot was covered with a plastic bag.

### Donald Danforth Plant Science Center

After a six week stratification in moist long fiber sphagnum moss (Luster Leaf Products Inc., USA) at 4 C, *Setaria* seeds were planted in four inch diameter (10 cm) white pots pre-filled with ~470 cm^3^ of Metro-Mix 360 soil (Hummert, USA) and 0.5 g of Osmocote Classic 14-14-14 fertilizer (Everris, USA). After planting, seeds were given 7 days to germinate in a Conviron growth chamber with long day photoperiod (16 h day/8 h night; light intensity 230 μmol/m^2^/s) at 31°C day/21°C night before being loaded onto the Bellweather Phenotyping System using a random block design. Plants were grown on the system for 25 days under long day photoperiod (16 h day/8 h night; light intensity 500 μmol/m^2^/s) cycling the temperature to match the photoperiod (31°C day/21°C night) and maintaining the relative humidity between 40 – 80 %. Weighing and watering of plants was performed between 2-4 times per day to maintain soil volumetric water content at either 40% full-capacity (FC) (drought) or 100% FC (well-watered) as determined by (32). Water limitation began 15 days after planting. Individual plants were imaged at 4 different angular rotations (0°, 90° 180°, 270°) every other day.

The effect of pot size was evaluated by growing parental lines of *Setaria viridis* (A10) and *Setaria italica* (B100) in four different size pots: 160 cm^3^ (T.O. Plastics, 715348C), 473 cm^3^ (Pöppelmann, VCC 10 F US), 950 cm^3^ (Stuewe & Sons, MT38) and 6033 cm^3^ (Nursery Supplies, C600) filled with Metro-Mix 360 soil (Sun Gro Horticulture). Seeds were given 7 days to germinate in a greenhouse and watered daily by the Donald Danforth Plant Science Center greenhouse staff. Plants were grown for 50 days under long day photoperiod (14 h day/ 10 h night) cycling the temperature to match photoperiod (27°C day/ 23°C night) and maintaining the relative humidity between 40-100%.

### Phenotyping plant height

In the field, plant height was measured as the distance from the plant base to the uppermost leaf collar on the culm. In 2013 this was measured by judging the average height of the canopy in a subplot. In 2014 height was directly measured repeatedly on three tagged plants in each subplot. In each field experiment, a direct culm height measurement was taken at the final destructive biomass harvest. In 2013 this was done on three representative plants from each subplot; in 2014 this was done on the same-tagged plants as the infield repeated height measurements.

In the 2013 Density experiment, height was measured at 19, 25, 31, 38, 45, 52, 59, and 67 days after seed sowing. Final biomass harvest began 87 days after seed sowing. In the 2013 Drought experiment, height was measured at 21, 29, 36, 43, 50, 59, and 67 days after seed sowing. Final biomass harvest began 92 days after seed sowing. In the 2014 Drought experiment, height was measured at 25, 33, 40, and 47 days after seed sowing. Final biomass harvest began 59 days after seed sowing. In the 2014 Density experiment, height was measured at 25 and 46 days after seed sowing. Final biomass harvest began 67 days after seed sowing.

At Carnegie, plant height was measured from the base of the plant to the tip of the panicle (at 48 days after seed sowing).

Five different measurement functions encoded within the PlantCV software package were used to estimate plant height in the images collected at the Bellweather Phenotyping Facility (32) (Figure S1). These include length of the plant object along the y-axis (extent_y), length of the plant object from the top of the pot to the maximal y-axis point identified as the plant object (height_above_bound), the distance from the top of the pot to the y-coordinate reported as the center of mass of the plant object (centroid_y), the distance from the top of the pot to the y-coordinate identified as the center of the ellipse that encompasses the plant object (ellipse_y) and an additional function designed to estimate the distance between the top of the pot and y-coordinate where average plant width is maximized (canopy_height). Canopy height is calculated by dividing the height_above_bound measure into 20 different equally sized scoring windows (divide the object into 5% intervals along the y-axis, scoring is based upon median width within each interval) and measuring the median plant width within each window. The function reports the average of y-coordinates that comprises the scoring window and median width within the scoring window where the object (plant) is the widest (canopy_width).

Centroid_y and ellipse_center_y were the most and least heritable measurements of plant height respectively (Figure S2a and Table S2). Height_above_bound and extent_y exhibited the largest proportion of variance attributed to water limitation (Figure S2b and Table S3). Summarization of height measurements derived from four rotational side view images was performed by calculating the mean, median, maximum or minimum of each distribution. Summarization method does not appear to influence the heritability of this these traits (Figure S3). All height metrics are reported as the average value estimated from all individual rotational images at each plant time point. Lengths of objects measured in pixels were converted to centimeters (cm) using the calibration reported by (32).

Plant height of parental lines grown in the greenhouse at the Donald Danforth Plant Science Center were measured as the distance from the base of the plant to most distant leaf collar on the culm using a tape measure. Measurements were recorded every other day during the workweek for a span of 50 days.

### Genotyping RILs

A de novo genetic map of the A10 × B100 *Setaria* RIL population was constructed through genotyping by sequencing (GBS) using the methods described in (34,35). Briefly, leaf tissue was collected from plants in the 2014 planting density experiment in addition and greenhouse trials. High molecular weight DNA was extracted using the CTAB method (36) and quantified using a Nanodrop 1000 Spectrophotometer [Thermo Scientific; Waltham, MA]. DNA was then double2 digested using the restriction enzymes *Pstl* and *MspI* and a unique barcode and common adaptors were ligated onto their respective fragment ends (37). After size selection and cleanup using Agencourt AMPure XP beads [Beckman Coulter; Brea, Ca], barcoded samples were pooled and submitted for sequencing (1002bp single end reads) using two Illumina Hi2Seq lanes [Illumina; San Diego, CA].

The resulting sequence reads were processed using the Tassel 3.0 GBS pipeline (38). Raw sequence reads were filtered to remove any tags that were not present 20 times at minimum within in each lane of sequencing and anchored to the *S. viridis* genome version 1.1 (Phytozome) using bowtie2 (39) with default parameters. Anchored SNP polymorphisms exhibiting an inbreeding coefficient less than 0.9 were removed and identical taxa within and between library preps (2 sequencing lanes) were called as heterozygous if the ratio between alleles in and between all identical taxa and/or library preps was less than 0.8. All polymorphisms categorized as heterozygous were called as missing values for the remainder of the analysis, yielding 17,612 SNPs.

The genetic map was generated using the R/qtl software (40). Genotype calls within the combined A10 taxa were compared to the *S. viridis* reference genome version 1.1 (Phytozome) and all missing and non-congruent polymorphisms were removed. The remaining sites were filtered to remove SNPs not present in > 90% of the unified taxa yielding 217 RILs and 3,295 markers. All polymorphisms were then recoded to reflect their parent of origin. Dubious genotype calls were identified by examining the recombination frequency between adjacent markers. Based upon manual inspection of histograms generated of recombination fraction between adjacent markers, all markers exhibiting a recombination frequency between adjacent markers greater than 36% (80 recombinations / 217 individuals) were considered to be erroneous and removed from the map. Next imputation of missing genotypes was performed at all sites that did not exhibit recombination between missing makers and duplicate markers were removed from the genetic map. Finally, the genetic distance between markers was calculated using the Kosambi mapping function assuming a genotype error rate of 0.005. Prior to QTL mapping, individuals missing more than 10% of their genotype calls are removed yielding a genetic map containing 208 individuals genotyped at 1,595 sites with 99.6% coverage. Based on pairwise similarity between identical individuals, the GBS map constructed here is highly consistent with other genetic maps of this population described in the literature (29,41).

### Partitioning trait variance, calculating heritability and estimating line values

Variance components corresponding to broad sense heritability and total variance explained were estimated using a mixed linear model using the R package lme4 (42). Broad sense heritability was calculated using two methods. Within an individual experiment, broad sense heritability on a line-estimate basis was calculated using the following formula:

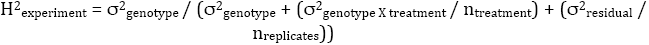

in which n_treatment_ is the harmonic mean of the number of treatment blocks in which each line was observed and n_replicates_ is the harmonic mean of number of replicates of each genotype in the experiment. Heritability within treatment blocks was calculated by fitting a linear model with genotype as the only explanatory factor within each treatment block.

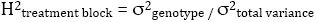

The proportion of variance attributed to genotype divided by total variance within each treatment block is reported as broad sense heritability within treatment.

Total variance explained was calculated by fitting a linear model including factors genotype, treatment, plot and genotype x treatment effects across all phenotypic values in all treatments. The proportion of variance that is incorporated into these factors divided by the total variance in the experiment is reported as total variance explained.

Best linear unbiased predictors (BLUPs) of plant height for each genotype were predicted from a linear mixed effect model independently within all individual time points of field drought and density experiments in 2013 and 2014 using lme4 (42). The following terms were included in the model:

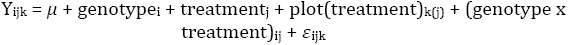

In which Y_ijk_ is the individual observation of plant height, *μ* is the overall mean, genotypei is the effect of the i_th_ genotype, treatment_j_ is the effect of the j_th_ treatment, plot(treatment)_k(j)_ is the effect of the k_th_ plot nested within the j_th_ treatment, and (genotype x treatment)_ij_ is a term describing the interaction between the i_th_ genotype and j_th_ treatment and *ε*_ijk_ is the random error term. As fixed effects we entered treatment and plot nested within treatment into the model. The terms genotype and genotype by treatment interactions were modeled as random effects.

Logistic regression was used to model the line values of plant height in experiments performed at the Bellweather Phenotyping Facility. Three types of logistic regression models considered (three2parameter logistic, fou^parameter logistic and Gompertz) and estimates from the model with the lowest Akaike information criterion (AIC) score were used. Modeling was performed using R functions described in (43).

### QTL analysis

QTL mapping was performed at each time point within treatment blocks and on the numerical difference, relative difference and trait ratio calculated between treatment blocks using functions encoded within the R/qtl and funqtl package (40,44). The functions were called by a set of custom python and R scripts (https://github.com/maxjfeldman/foxy_qtl_pipeline). Two complimentary analysis methods were utilized. First, a single QTL model genome scan using Haley-Knott regression was performed to identify QTL exhibiting LOD score peaks greater than a permutation based significance threshold (α = 0.05, n = 1000). Next, a stepwise forward/backward selection procedure was used to identify an additive, multiple QTL model based upon maximization of penalized LOD score. Both procedures were performed at each time point, within treatment blocks and on the numerical difference relative difference and trait ratio calculated between phenotypic values measured in treatment blocks at each time point. QTL associated with difference or ratio composite traits may identify loci associated with genotype by environment interaction (45).

The function-valued approach described by Kwak et al., 2016 (44) was used to identify QTL associated with the average (SLOD) and maximum (MLOD) score at each locus throughout the experiment (44). The genotypic mean height within treatments was estimated using a logistic function and the QTL significance threshold was determined based upon permutation-based likelihood of observing the empirical SLOD or MLOD test statistic. Across all grow outs, separate; independent linkage mapping analysis performed at each time point identified a larger number of QTL locations relative to similar function valued analysis based on the SLOD and MLOD statistics calculated at each individual marker throughout the experimental time course.

After refinement of QTL position estimates, the significance of fit for the full multiple QTL model was assessed using type III analysis of variance (ANOVA). The contribution of individual loci was assessed using drop-one-term, type III ANOVA. The proportion of variance explained by in addition to the allelic effect size of each locus were determined by comparing the fit of the full model to a submodel with one of the terms removed. All putative protein coding genes (*Setaria viridis* genome version 1.1) found within a 1.5-logarithm of the odds (LOD) confidence interval were reported for each QTL.

## Results

### Evaluating different measurements of plant height

Height in the field was measured using two different methods of approximation. An analysis of variance indicates that on average, measuring the exact height to the collar of the uppermost leaf on the culm of a subset of three individuals within a plot (as performed in 2014) was approximately 5.8% more heritable than attempting to estimate the average height to the collar of the uppermost leaf on the culm of all plants within the plot (as performed in 2013) (Table S1). An increase in heritability between years was not observed for flowering time between the 2013 and 2014 field seasons, suggesting that the increase in heritability was not due to differences in environment or management (Table S1).

Five different automated methods of measuring plant height (height_above_bound, extent_y, centroid_y, ellipse_center_y, canopy_height - see definitions in methods) from 2-D images captured at the Bellweather Phenotyping Facility were evaluated (Figure S1). Among these measurements, height_above_bound, extent_y and centroid_y were highly correlated (R^2^ > 0.97) (Table S2) and all three were highly heritable on average (h^2^ > 0.85) and showed clear effects of water limitation (Figure S2, Table S3 and Table S4). Summary statistic does not greatly influence the heritability of height type measurements (Figure S3). The mean value of height_above_bound based upon four rotational images was the primary metric used for the remainder of this study as it is intuitively simple to understand, highly heritable, strongly correlated with other height metrics, sensitive to treatment effects and previously validated against manual measurements (32).

### Variation in plant height

Height of individuals within the *S. viridis* (A10) X *S. italica* (B100) RIL population was evaluated under very diverse growing conditions. Variation resulted from different water availability and planting density treatments. In addition, day length and temperature varied between plant2outs at three locations (Illinois field site, DDPSC growth chamber, Carnegie growth chamber) and two different planting dates each year at the Illinois field site. Although the mean and variance of final height differed significantly between grow outs (Figure 1a), the coefficient of variation remained relatively constant across all replicated experiments and treatments (Table 1 and Table S5). The proportion of variance attributed to experimental factors (genotype versus treatment versus error) also varied between trials (Figure 1b) and throughout time in a given experiment (Figure 1c and Table S4). Across all experiments the mapping population size averaged ~160 individuals with a minimum count of 146 (Table 1). Final height estimated near the end of each these experiments averaged 38 cm but every experiment exhibited at least a 20 cm difference between the 5^th^ and 95^th^ percentile (Table 1). Modest transgressive segregation was apparent across all experimental trials (Figure 1a). At the end of all eight field grow outs, the B100 parental line was taller than the A10 parent, whereas in three out of four grow outs conducted in artificial environments at Carnegie and Bellweather, A10 was taller (Figure 1a).

**Fig 1.**
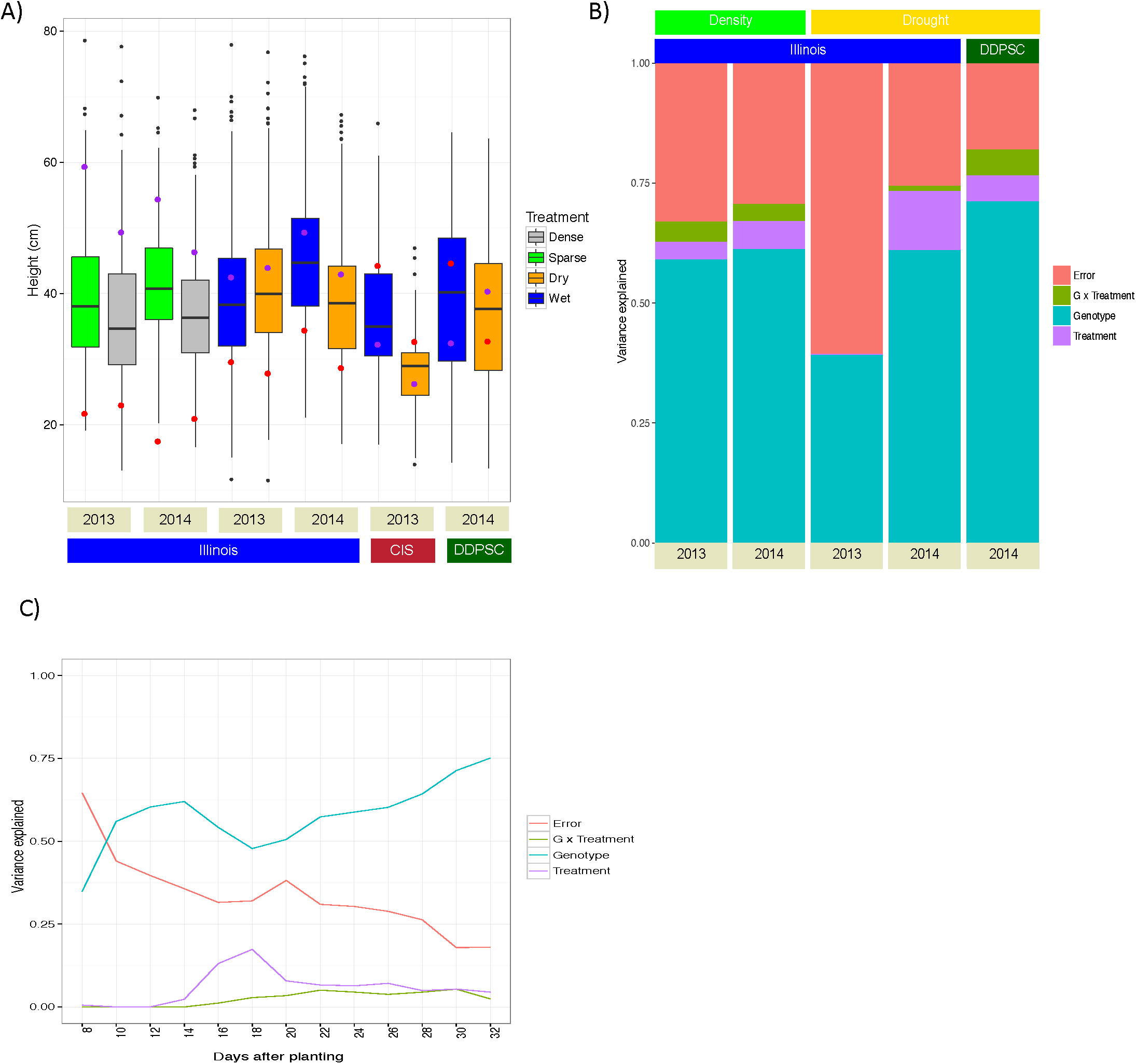
A large proportion of variance can be attributed to genotype and treatment factors, the effect of which changes throughout time and across trial locations. Field grow outs were performed at the University of Illinois Champaign-Urbana (Illinois), whereas green house experiments were performed at the Carnegie Institute for Science (CIS) and the Donald Danforth Plant Science Center (DDPSC). Both water availability and planting density experiments were conducted at Illinois in 2013 and 2014. A) Boxplot summaries of plant heights at the end of each experiment. Red points indicate the mean height of the A10 parental line and purple points denote the mean height of the B100 parental line. B) The proportion of variance that can be attributed to experimental factors at the end of each study that included replication. C) The proportion of variance that can be attributed to experimental factors throughout the Bellweather 2014 grow out.

**Table 1.** Summary of experimental metrics from field and controlled environments. Time point corresponds days after planting, plant height is abbreviated as PHT and coefficient of variation is abbreviated as CV.

| <u>Location</u> | <u>Environment</u> | <u>Year</u> | <u>Time point</u> | <u>Treatment</u> | <u>Summarization method</u> | <u>Genotypes planted</u> | <u>Mapping pop. size</u> | <u>Replication</u> | <u>PHT 5%</u> | <u>PHT 50%</u> | <u>PHT 95%</u> | <u>Mean</u> | <u>Variance</u> | <u>CV</u> | <u>Correlation with panicle emergence</u> |
| --- | --- | --- | --- | --- | --- | --- | --- | --- | --- | --- | --- | --- | --- | --- | --- |
| Illinois | Field | 2013 | 92 | Dry | BLUP | 183 | 159 | 2.90 | 25.47 | 39.95 | 56.80 | 40.50 | 98.93 | 0.33 | 0.30 |
| Illinois | Field | 2013 | 92 | Wet | BLUP | 170 | 159 | 2.99 | 24.20 | 38.29 | 56.20 | 39.16 | 98.19 | 0.37 | 0.31 |
| Illinois | Field | 2013 | 87 | Dense | BLUP | 183 | 169 | 2.40 | 21.04 | 34.68 | 51.67 | 36.03 | 96.63 | 0.34 | 0.69 |
| Illinois | Field | 2013 | 87 | Sparse | BLUP | 182 | 168 | 2.96 | 22.15 | 38.06 | 59.70 | 39.19 | 118.10 | 0.34 | 0.75 |
| Illinois | Field | 2014 | 59 | Dry | BLUP | 157 | 146 | 3.00 | 25.42 | 38.53 | 57.28 | 38.90 | 88.61 | 0.28 | 0.37 |
| Illinois | Field | 2014 | 59 | Wet | BLUP | 157 | 146 | 3.00 | 30.41 | 44.70 | 64.79 | 45.28 | 102.31 | 0.28 | 0.37 |
| Illinois | Field | 2014 | 67 | Dense | BLUP | 188 | 174 | 1.97 | 21.87 | 36.34 | 53.36 | 36.76 | 81.59 | 0.31 | 0.49 |
| Illinois | Field | 2014 | 67 | Sparse | BLUP | 187 | 172 | 1.93 | 27.13 | 40.76 | 56.44 | 41.59 | 82.06 | 0.29 | 0.42 |
| Stanford | Growth chamber | 2013 | NA | Dry | None | 160 | 149 | 1.00 | 18.00 | 29.00 | 38.45 | 28.31 | 40.05 | 0.22 | NA |
| Stanford | Growth chamber | 2013 | NA | Wet | None | 166 | 155 | 1.00 | 23.43 | 35.00 | 51.08 | 36.65 | 82.21 | 0.25 | NA |
| Danforth | Growth chamber | 2014 | 30 | Dry | Logistic regression | 187 | 157 | 2.91 | 18.85 | 37.64 | 58.36 | 37.44 | 134.69 | 0.31 | NA |
| Danforth | Growth chamber | 2014 | 30 | Wet | Logistic regression | 185 | 167 | 2.89 | 20.60 | 40.21 | 59.72 | 39.40 | 146.29 | 0.31 | NA |
| AVERAGE | AVERAGE | NA | NA | NA | NA | 175.42 | 160.08 | 2.41 | 23.22 | 37.76 | 55.32 | 38.27 | 97.47 | 0.30 | 0.46 |

Within each trial and treatment, the heritability of final plant height exceeded 0.84 (Table S1). The mean heritability of plant height in this population across all time points was 0.82 and tended to be higher in controlled environments (average H^2^ = 0.86 at Bellweather and Carnegie, average H^2^ = 0.83 in field at Illinois). Generally, the proportion of height variance attributed to genotype increased as the plant approached maturity (Figure 1c and Table S6). The effects of water limitation and planting density treatments did not dramatically influence the heritability of plant height in this study (Table S1).

The block structure of the field experimental design and the frequent repeated measurements on the Bellweather phenotyping system enabled modeling of the underlying trait phenotypes. Best linear unbiased estimated line values of height within individual experiment and time points were used to account for variation attributed to plot and treatment effects in field studies. Logistic regression was used estimate the expected value of repeated temporal measurements of height collected at the Bellweather Phenotyping Facility.

### Variation of plant height in response to growth environment and experimental treatment

Water limitation experiments were conducted in all three locations; each of which differed by genotypes planted, statistical power and the severity of treatment administered in addition to other latent variables inherent to growth location and experimental design. The effect of these factors was assessed at a single time point representative of final plant size using type III analysis of variance (46). Within drought trials, genotype had the strongest effect on height, followed by trial location and treatment effect nested within location (Table S7). Similar to the correlation of height and days to anthesis in maize (13), a significant positive correlation between final height and day of panicle emergence was observed in each field grow out (Table 1), suggesting that different planting dates, with their different day lengths and temperatures could explain some of the trial location variation.

The water limitation treatment varied widely between experiments, simulating several possible scenarios plants could experience. Field trials at Illinois and the controlled growth trial at Carnegie were designed as progressive drought experiments, where after an initial growth period, water is withheld. In the Bellweather phenotyping experiment, a reduced water availability set point was maintained after an initial dry down. The controlled growth experiments all showed a significant effect of drought, as did the 2014 field experiment (despite an end of season flood). In retrospect, water was not withheld early enough in the 2013 field experiment to detect a significant drought effect on height although soil probes showed a substantial difference in water availability within 3 weeks (Table S8). The effect of withholding water completely soon after germination in the Carnegie experiment made the observed height effect the most pronounced of any grow out observed (Figure S5). Height appears less responsive to planting density than water limitation, with most of the variance attributed to genotypic and plot nested within treatment followed by the variance attributed to year and lastly planting density (Table S7). Nonetheless, in both 2013 and 2014, planting density had a significant effect on height by the end of the experiment.

Using the rank order of lines by phenotype as a proxy for how similar the experiments are to each other, the effect of experimental location is apparent, as well as the difference between controlled environment and the field. (Figure 2 and Figure S4). The well-watered treatment at Bellweather, the drought treatment at Carnegie and the 2013 field density experiment are the most divergent environments encountered within this study (Figure 2).

**Fig 2.**
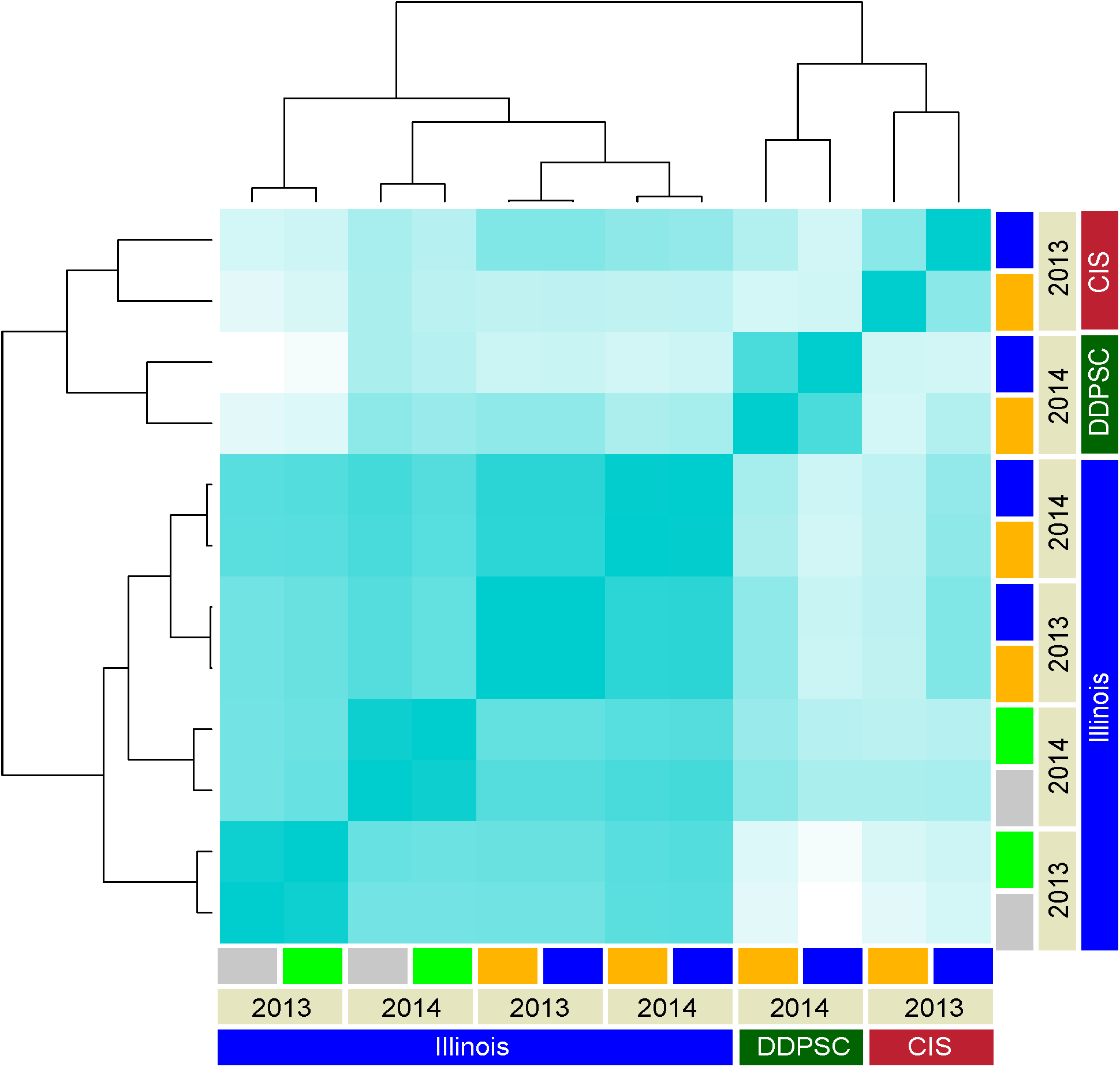
Rank order correlation of plant height measured at the end of each experiment. Grow outs of the A10 X B100 RIL population were performed in three different locations including the University of Illinois Champaign-Urbana, Carnegie Institute for Science (CIS) and Donald Danforth Plant Science Center (DDPSC). Treatment block is indicated by color bar at the edge of the dendrogram (blue corresponds to wet, orange represents to dry, green indicates sparse and grey denotes dense planting). The intensity of color shading is proportional to the magnitude of Pearson’s correlation coefficient (0.0 is white whereas 1.0 is blue).

### Genetic map construction for the *Setaria* A10 X B100 RIL population

Two previous genetic maps have been constructed for this population (29,41) but neither included all of the lines used in this study. We constructed a *de novo* genetic map and genotyped all phenotyped RILs to verify the identity of genetic stocks. DNA samples were prepared from 183 field grown and 34 greenhouse grown RIL lines (in addition to the two parental genotypes) and GBS libraries were prepared and sequenced (34,35). After alignment to the recently completed *S. viridis* genome (Phytozome v1.1) using bowtie2 (39) and filtering to select SNPs that match the sequenced parent samples and removal of duplicates, the final genetic map consists of 217 individuals, spanning 1,088 cM, calculated using 1,595 markers (Table S9). After imputation, the average spacing between markers is 0.7 cM with the greatest distance between markers being 14.4 cM (Table S9). The map size, coverage and identity of individuals in this RIL population are largely identical between map builds (29,41).

### Architecture of height QTL in different environments

We implemented a standard stepwise forward/backward selection procedure to identify the optimal, additive, multiple QTL model based upon penalized LOD score (40). Mapping was performed on all traits, at each time point within treatment blocks and on the numerical difference, relative difference and trait ratio between phenotypic values by treatment blocks. This procedure identified a total of 153 individual SNPs associated with height across all time points 1. 0 is blue). and treatments within the 12 experimental grow outs (Figure 3 and Table S10). Many of the individually identified SNP positions group into clusters of linked loci that are likely representative of a single QTL location. Groups of closely linked QTL markers (10 cM radius) were collapsed into the most frequently observed SNP marker in each cluster, resulting in 37 unique QTLs (Table S11 and Figure S6). A substantial proportion of these QTL were observed at least once in three (17/37) or four (12/37) treatment groups, suggesting that the genetic architecture of height is similar across all experimental treatments contrasts considered in this study (Table S11).

**Fig 3.**
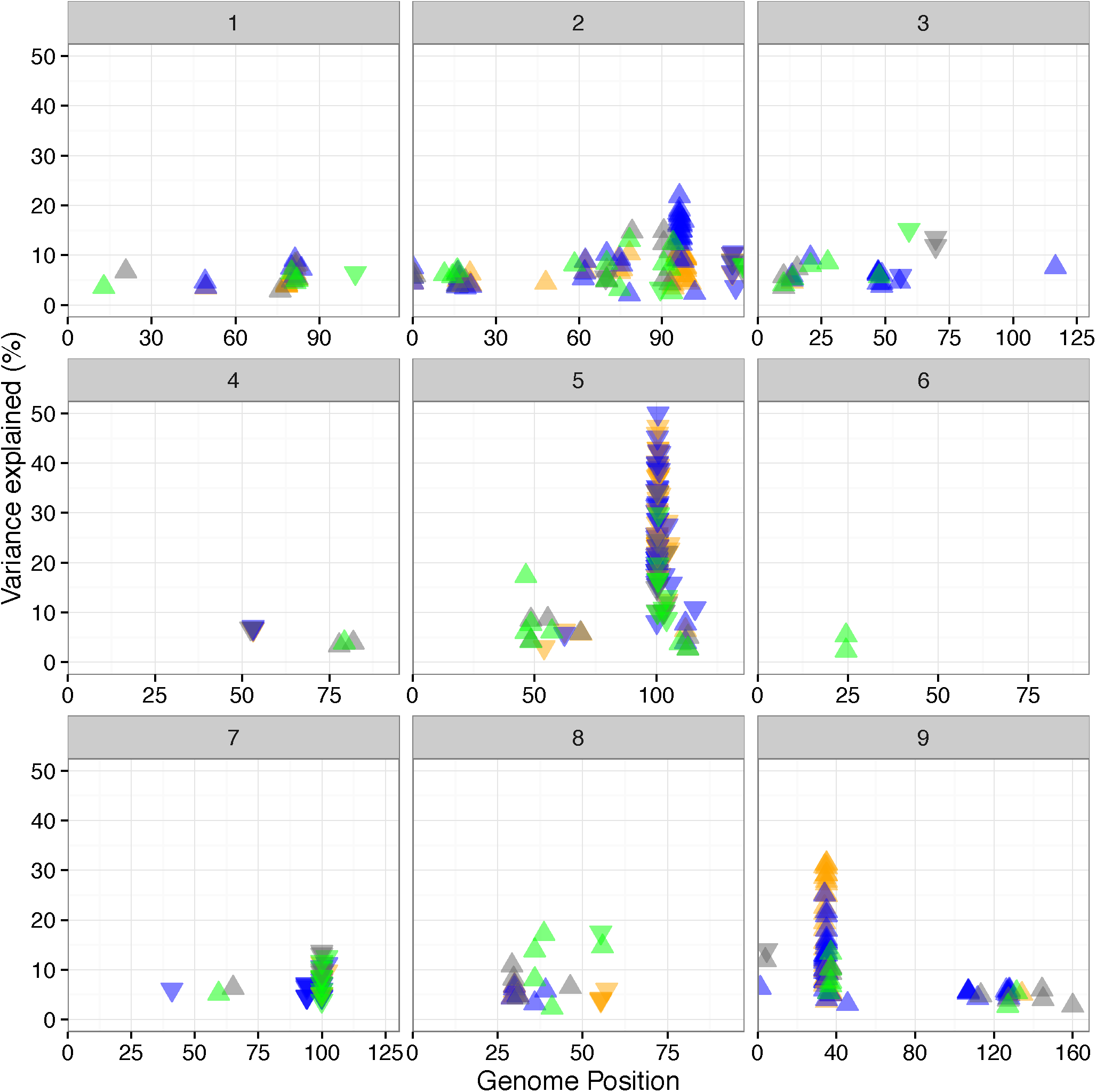
One hundred and fifty three unique QTL positions were identified across all experiments. Each box corresponds to an individual chromosome, where the values along the x-axis are chromosome position and values along the y-axis denote the proportion of genetic variance explained by the QTL. Each triangle represents a single QTL detected, where the color indicates the treatment condition the QTL was identified in (blue represents wet, orange corresponds to dry, green indicates sparse planting density whereas grey denotes dense planting) and direction of the arrow corresponds the directional effect of the B100 parental allele.

A static QTL model including only the 12 QTL locations identified at least once in all treatment groups was fit at a time point representative of final plant size in each experiment. Across all experimental grow outs these 12 QTL explained between 25 - 64% of the additive genetic variance and 53% on average (Figure 4). Across all experimental time points in this study the three loci that explain the greatest proportion of variance in any single time point are included within this set of 12 QTL. The most frequently observed QTL (marker: S5_41999990) was identified in each of the 12 experiments and explained the largest percentage of additive genetic variance (mean of 19%). The B100 allele at this QTL position is associated with reduced plant height. The individual contribution of the other 11 QTL at this representative time point on average, did not explain more than 7% of the additive genetic variance alone, but cumulatively these 11 QTL account for greater than 33%. The B100 allele at nine of these twelve positions are associated with increased plant height.

**Fig 4.**
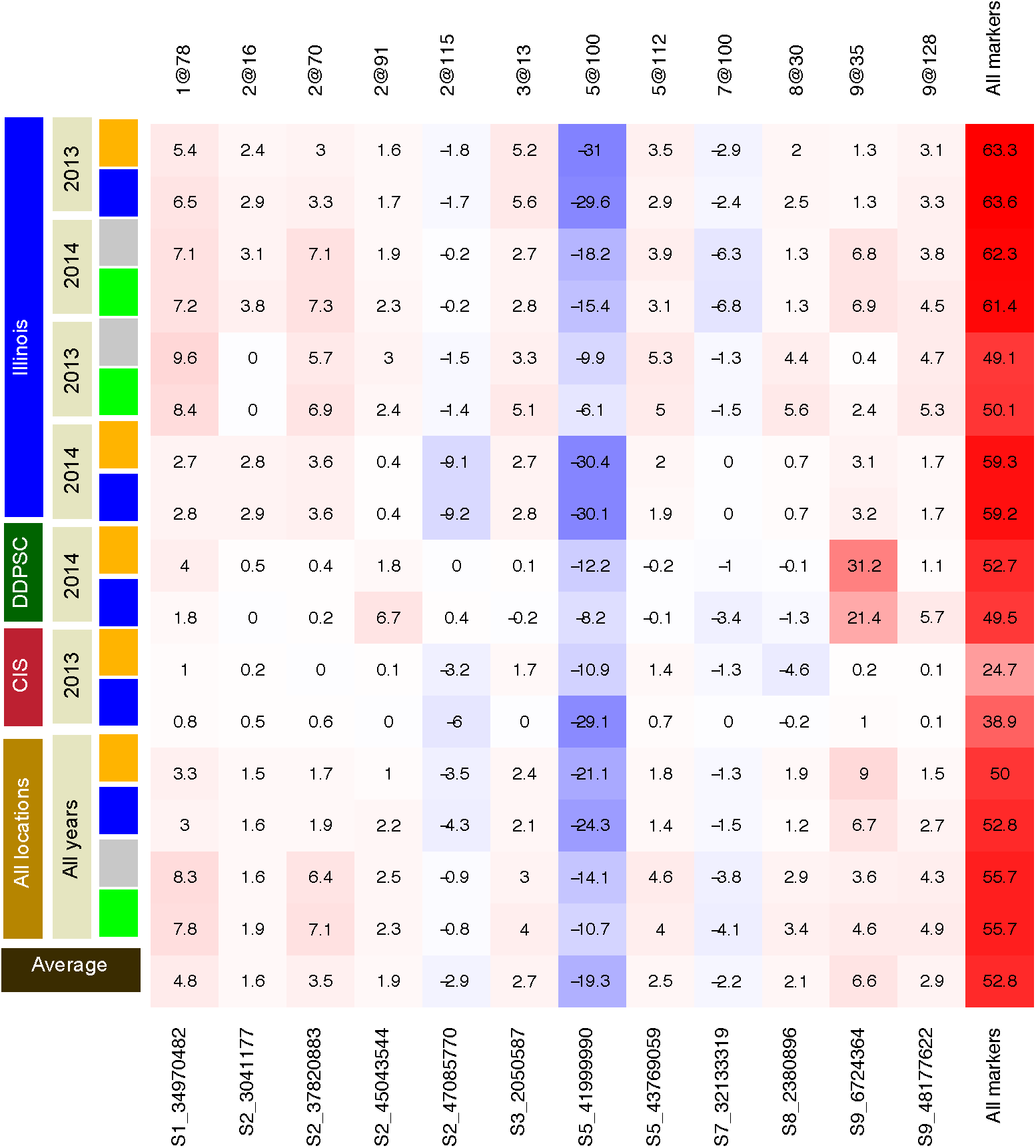
Twelve QTL were identified across all experimental treatment groups. Cells shaded red indicate a positive effect from inheriting the B100 parental allele whereas cells shaded in blue indicate a positive effect of inheriting the allele derived from the A10 parent. Values within the cells and intensity of coloration report the percentage of additive genetic variance explained.

Across all field trials performed, the number of QTL shared between treatment blocks and across years is greater than the number of QTL unique to any treatment block across growth years or both treatment blocks within a single growth year (Figure 5 and Figure S7). No overlap between QTL detected exclusively in well-watered or water-limited conditions was observed between any of the experiments (Figure 5a and Figure S7). Within planting density experiments a single QTL (marker: S9_52254840) was identified uniquely within the high planting density treatment block in both years (Figure 5b and Figure S7). A majority (13/16) of the QTL detected in controlled environments were also detected in field environments (Figure 5c).

**Fig 5.**
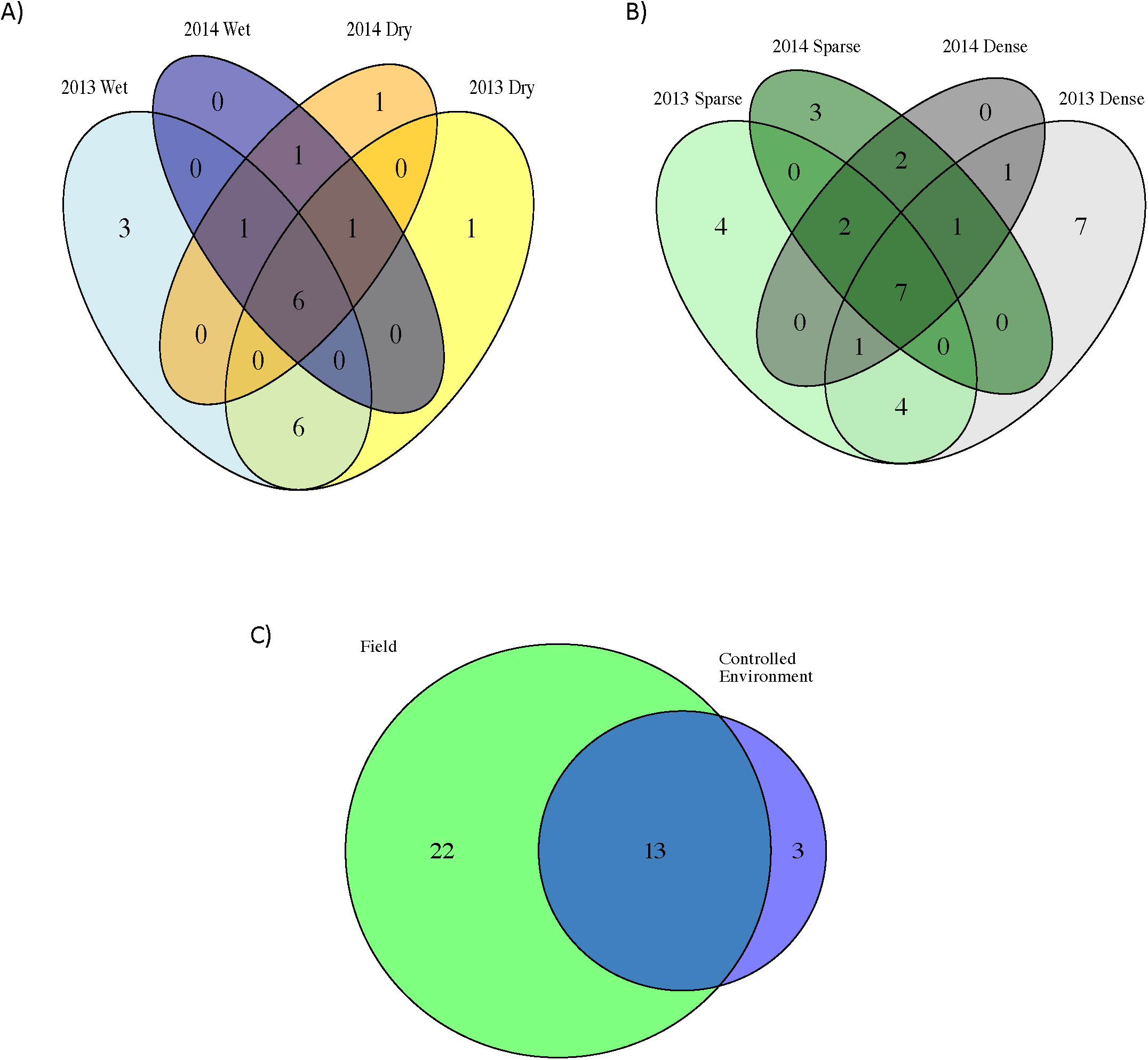
QTL associated with plant height are shared across experimental contrasts. A) No QTL unique to either treatment block were observed across both water limitation field trials in 2013 and 2014. B) In planting density experiments, one QTL was identified uniquely within the high density treatment block in field trials in both years. C) Almost all QTL identified in controlled environments are also identified in field environments.

The results from QTL scans performed on the numerical difference, relative difference and trait ratio between plant heights measured between treatment blocks at all time points largely overlapped with the results from the QTL analysis performed on measured value of plant height within treatments (Figure S8 and Table S12). In total, 18 of the 29 unique QTL associated with these numerical metrics of difference in plant height between treatment blocks were also identified during our QTL analysis of plant height (Figure S8 and Table S12). In contrast to the QTL locations identified for plant height, most of which were found in multiple experiments, of the 11 difference QTLs that do not overlap 10 were only found in one experiment (Figure S8 and Table S12). While there are genetic loci that appear to be responsive to different environments, the majority of the environmentally responsive loci are due to relative differences in the timing of the loci growth affect between treatments.

### Temporal architecture of plant height QTL

Height is a dynamic trait that changes at different rates throughout the plant life cycle. To determine which growth stage the identified loci influence, we examined the best fit multip^QTL model produced at each separate independent time point and performed an analysis using the average LOD (SLOD) and maximum LOD (MLOD) function-valued approaches as described by (44,47). Across all grow outs, linkage mapping analysis performed independently at each time point identified a larger number of QTL locations relative to the function-valued analysis. The positional locations of major QTL detected using each of these methods varied slightly, but in all cases it is likely that the same QTL are identified. To simplify discussion, we will refer to these QTL regions using notation that describes the approximate location of the QTL on each chromosome (QTL detected on chromosome 5 at position 99.1 cM and at position 105.2 will be reported as 5@100 for example).

The relative significance of each QTL location changes substantially throughout the life cycle of plant (Figure 6 and Figure S9). In all experimental grow outs, plant height at early stages of development is strongly associated with a single major QTL located at 5@ 100. In the dataset with the highest temporal resolution (Bellweather), the proportional contribution of this QTL diminishes over time while the contribution of other QTL located at 2@91 and 9@35 increase (Figure 7a). The proportion of variance explained by 5@100 diminished late in plant development in all field grow outs (Figure S10). However, height measurements started later and continued longer through plant development under field conditions. This revealed that the proportion of variance explained by 5@100 peaked between 30-40 days after sowing, before starting to decline. This is in contrast to its declining role throughout the growth period (8-32 days) under controlled environment conditions. Three out of four field studies also detected 9@35, and again the proportion of variance explained by this QTL diminished at late developmental stages only assessed in the field (Figures_S10). The magnitude of allelic effects contributed by individual loci generally increase through time in all growth environments (Figure 7b and Figure S11) highlighting the cumulative nature of developmental traits.

**Fig 6.**
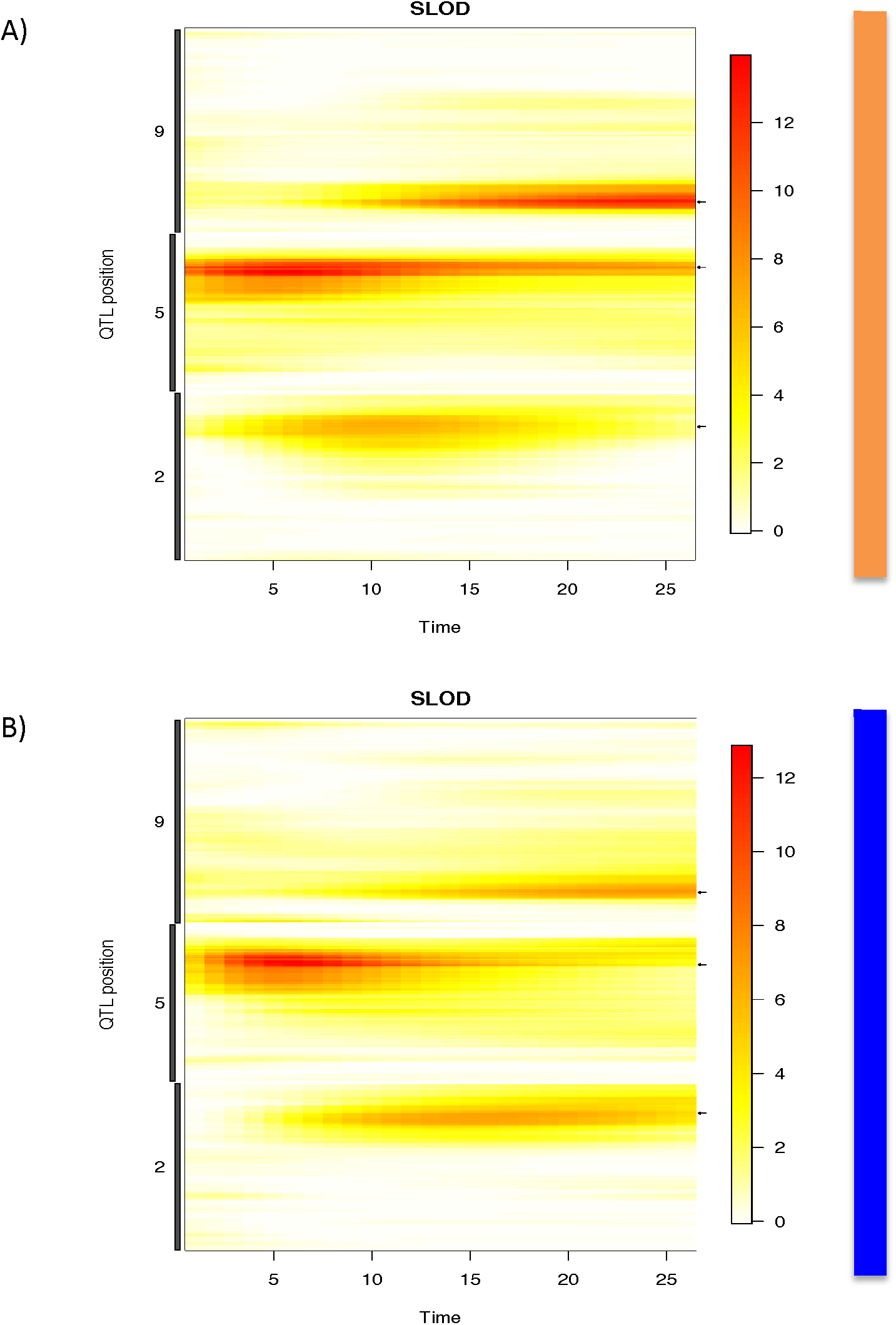
Three major QTLs account for the most significant proportion of variance across all time points during the Bellweather experiment. The significance of association at each QTL location is plotted over the course of the experiment. Coloring of the heat map is reflective of LOD score at genome location (cM) at a given time point. Increased red coloring is indicative of higher LOD score. A) Plot of LOD score at loci that significantly influence plant height in the well-watered treatment block of the Bellweather experiments as calculated using the SLOD test statistic. B) Plot of LOD score at loci that significantly influence plant height in the water-limited treatment block of the Bellweather experiments as calculated using the SLOD test statistic.

**Fig 7.**
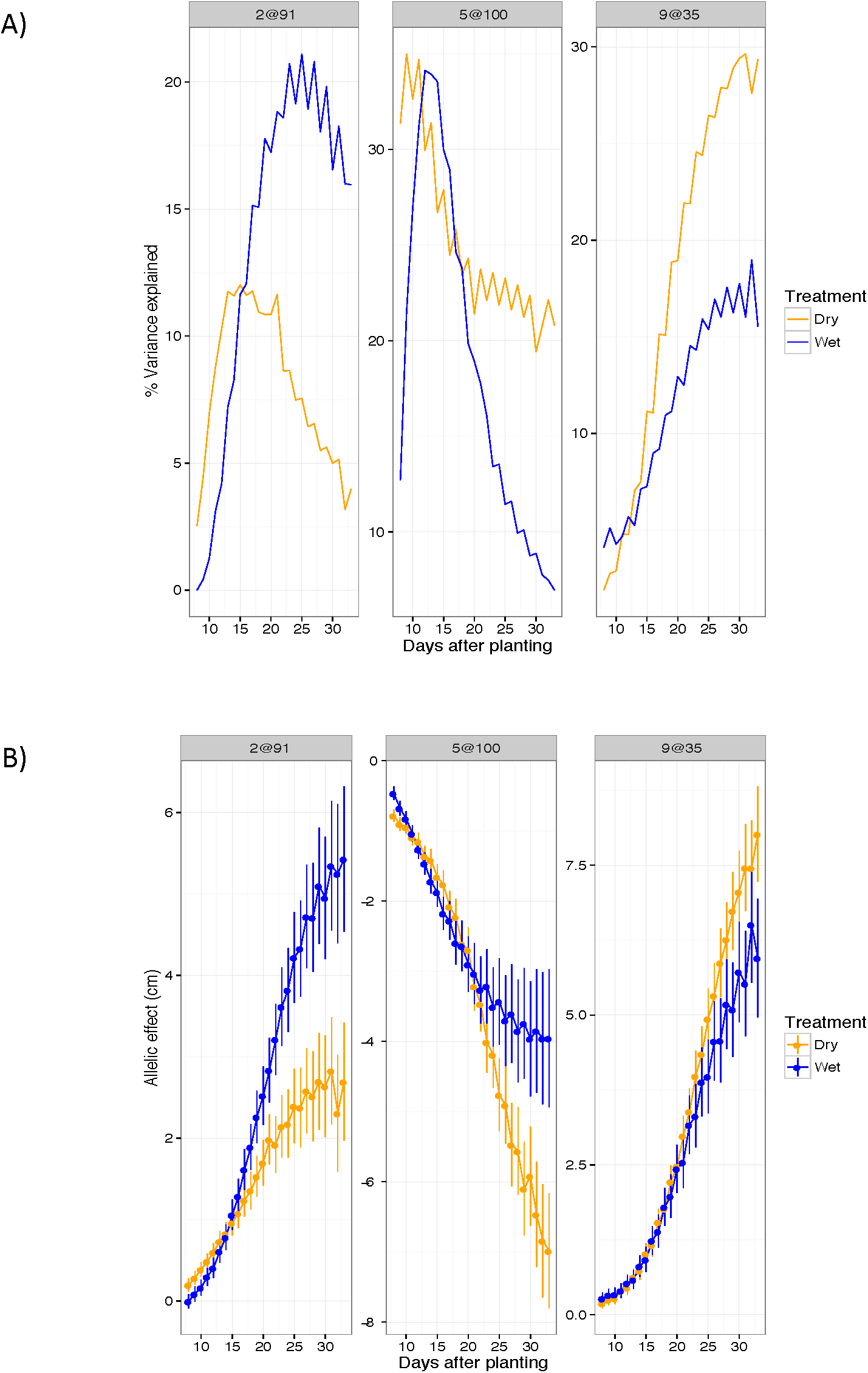
The temporal influence of individual loci influences effect sizes throughout development. QTL location is indicated with the following notation (chromosome@position). A) The proportion of variance attributed to the QTL at position at 2@91 and 9@35 is increases throughout the plant life cycle whereas the influence of the major QTL at position 5@100 decreases as the plant approaches maturity. B) The effect sizes of each locus are cumulative through out development. Although by the end of the experiment the proportion of variance explained by the locus at 5@100 is very low, the effect size continues to grow in magnitude.

Although the identity of the major loci that influence plant height in this population is relatively unresponsive to treatment, the proportional contribution of each locus throughout time appears responsive to water treatment (Figure 7). Water-limited plants exhibit slightly decreased average and maximal rates of growth (Figure 8). In addition, the day at which maximal growth rate is achieved is significantly delayed in water-limited plants (Figure 8). QTLs associated with maximal and daily growth rate are essentially identical to those detected for height, although the time at which these QTL exhibit the greatest influence appears shifted toward earlier time points in the experiment (Figure S12). No significant QTL were identified for either the day at which maximal growth rate occurs or the difference of this trait between treatments.

**Fig 8.**
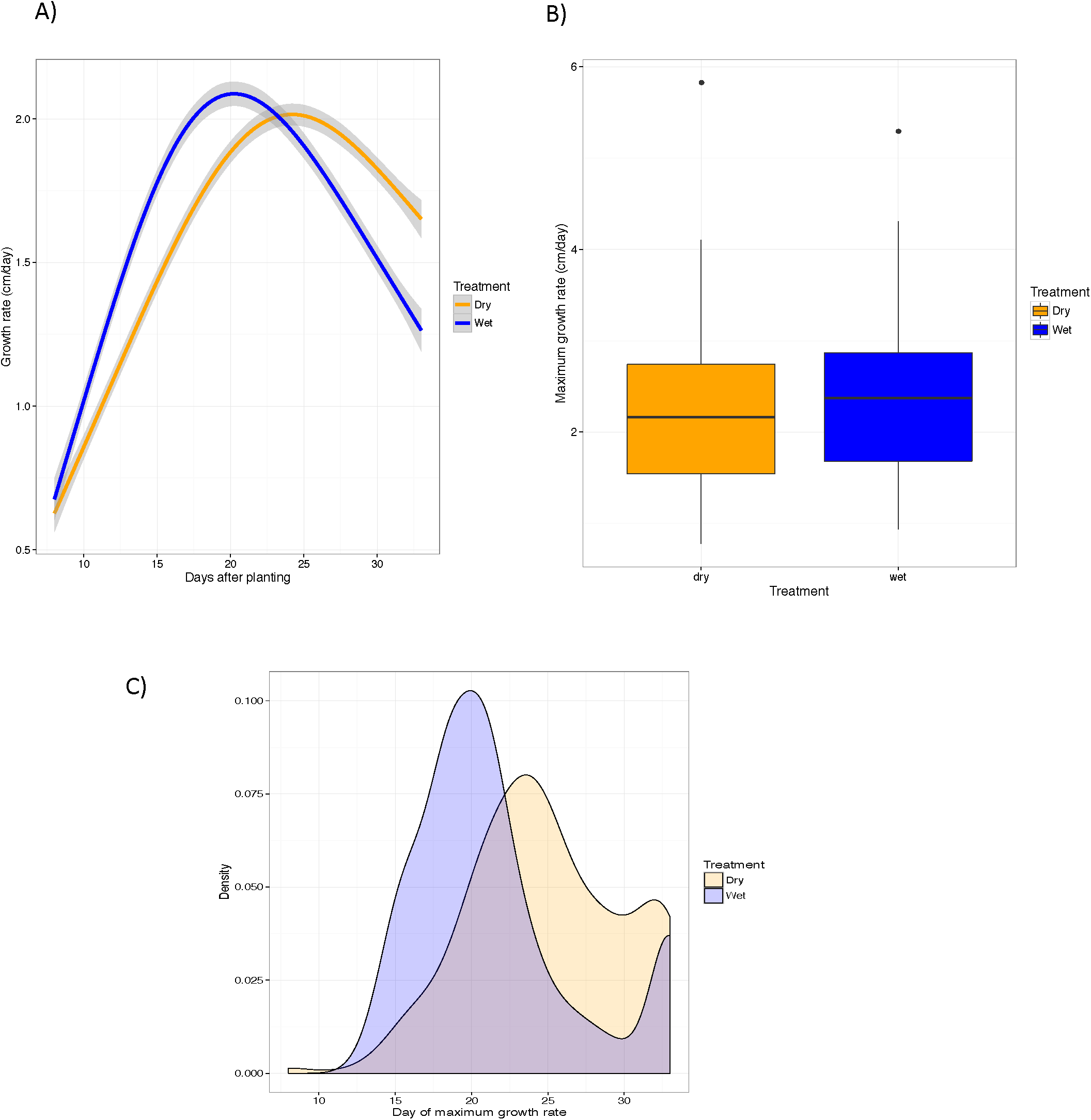
A delay in the developmental program that controls vertical growth is likely more influential on final plant height observed rather than maximal growth rate. A) Vertical growth rate plotted through time indicate rates of vertical growth vary dramatically throughout development. B) No significant differences (*α* =0.05) in the rate of maximal vertical growth was not observed between water level treatment blocks. C) Plants within the well-watered treatment block achieve their maximal vertical growth rate significantly earlier than their water limited counterparts.

### A temporal multi locus model of *Setaria* plant height

The SLOD based function-valued QTL analysis of our highest resolution dataset identified a simple, additive, QTL model comprised of loci on chromosomes 2, 5 and 9 (2@91, 5@100 and 9@35) which provides a framework to link many of the experiments. The same causative QTL loci were identified in the wet and dry treatment blocks of the grow outs at Bellweather. Early in plant development, individuals that inherit the B100 allele at position 5@100 will be shorter than those that inherit the A10 allele (Figure 9). The effects of this locus are then counterbalanced by the influence of the QTL at position 2@91 and later by 9@35 where the B100 allele increases plant height at maturity. If the RIL genotypes are divided into four genotypic classes based on these three loci: all A10 (AAA), all B100 (BBB) and combinations where the 5@100 loci is discordant (ABA and BAB), the effect of these loci is clear. The BAB class is the tallest and the ABA class is the shortest, throughout the growth cycle. The AAA and BBB classes, which include the parent genotypes, however, have an inflection point between the early growth, where AAA is initially taller and the later stages where the BBB class becomes and remains taller for the remainder of the experiment (Figure 9). In the well-watered treatment block of the Bellwether grow out, individuals that have inherited the AAA loci exhibit the largest initial growth rate of all genotypes but are over taken by both the BBB and BAB within 13 days after planting (Figure 9c). Individuals that inherit the BBB allele combination maintain the highest rate of vertical growth at the end of this experiment in the well-watered treatment block (Figure 9c).

**Fig 9.**
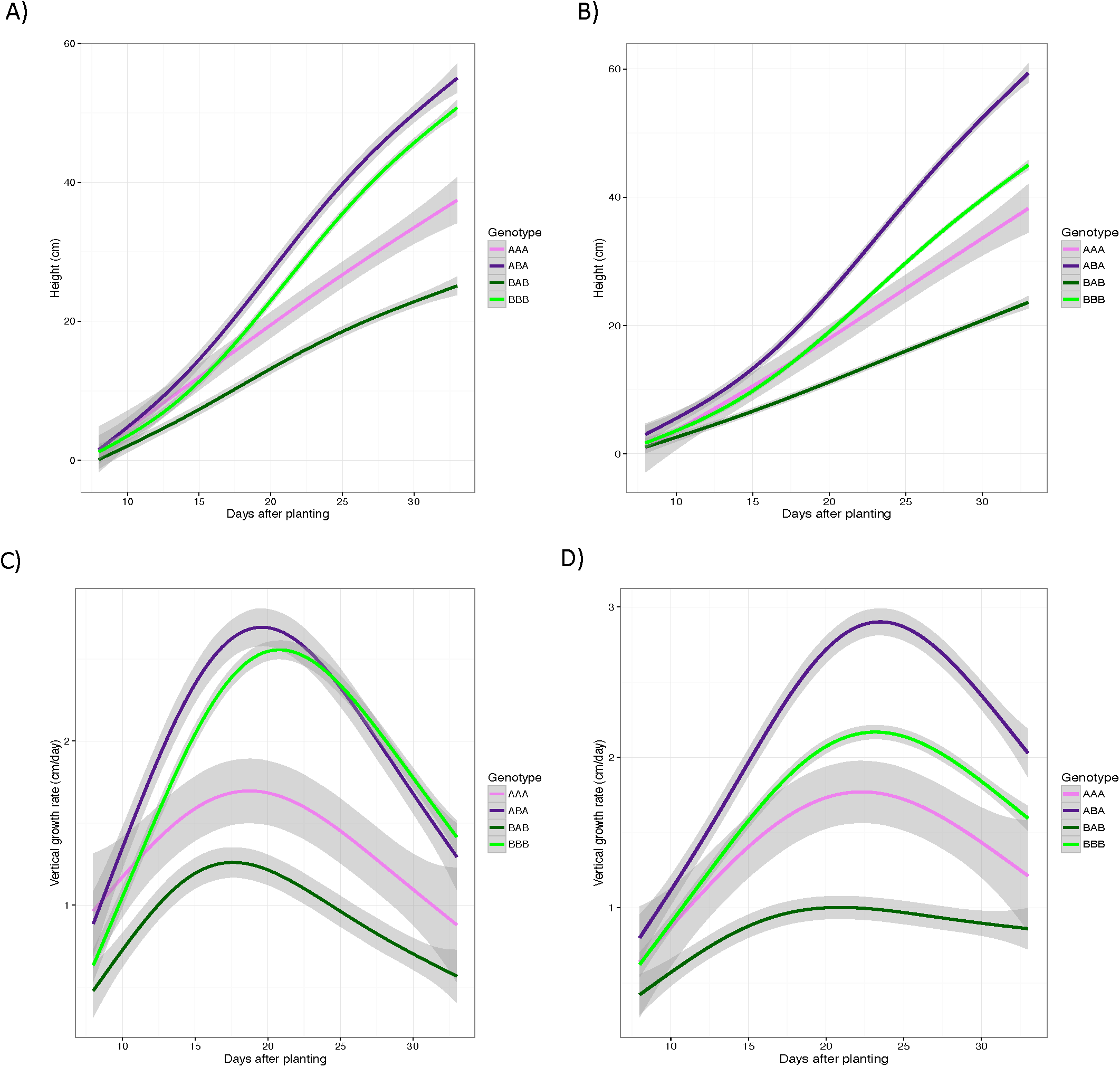
Plants that inherit A10 alleles at 5@100 exhibit greater plant height initially, whereas those that inherit alleles from the B100 parent at 2@91 and 9@35 generally achieve greater final heights. Genotypes are reported as three character string where individual characters represent the parental origin of the allele at locus 2@91, 5@100 and 9@35 respectively. A) The plant height of individuals that that exhibit parental genotypes (AAA and BBB) and those that are segregating at the locus at chromosome 5 (BAB and ABA) in well-watered conditions. B) The plant height of individuals that that exhibit parental genotypes (AAA and BBB) and those that are segregating at the locus at chromosome 5 (BAB and ABA) in water-limited conditions. C) The vertical growth rate of individuals that that exhibit parental genotypes (AAA and BBB) and those that are segregating at the locus at chromosome 5 (BAB and ABA) in well-watered conditions. D) The vertical growth rate of individuals that that exhibit parental genotypes (AAA and BBB) and those that are segregating at the locus at chromosome 5 (BAB and ABA) in water?limited conditions.

The 5@100 QTL is found in all of the experiments with the A10 contributing the tall allele. The 2@91 and 9@35 QTLs are present in many other experiments, but there are also several additional QTLs with large effect and B100 contributing the taller allele. In most of the experiments, a fairly simple model with the 5@100 QTL driving early growth and several B100 tall QTLs taking over later explains the observed variation (Figure 10). The exceptions are three field grow outs (2013 Dry, 2014 Dry/Wet) where A10 high QTLs drive early growth, but the B100 QTLs do not appear to be the major loci influencing growth at later developmental time points. Closer inspection of the variance explained by the QTLs in these experiments reveals that at later time points, the percent variance explained by the QTLs is low, despite the trait heritability remaining high (Figure S13). Running the QTL analysis with a lower permutation threshold cutoff (α = 0.25) reveals the presence of several additional small effect QTLs with B100 tall alleles. Given the limitations of the population size, it is reasonable to predict that there may be more small effect QTLs that could be contributing to make B100 taller in this experiment. This chronological genetic model of plant height may in part explain the differences in plant height observed when parental lines are grown in controlled environments versus field settings.

**Fig 10.**
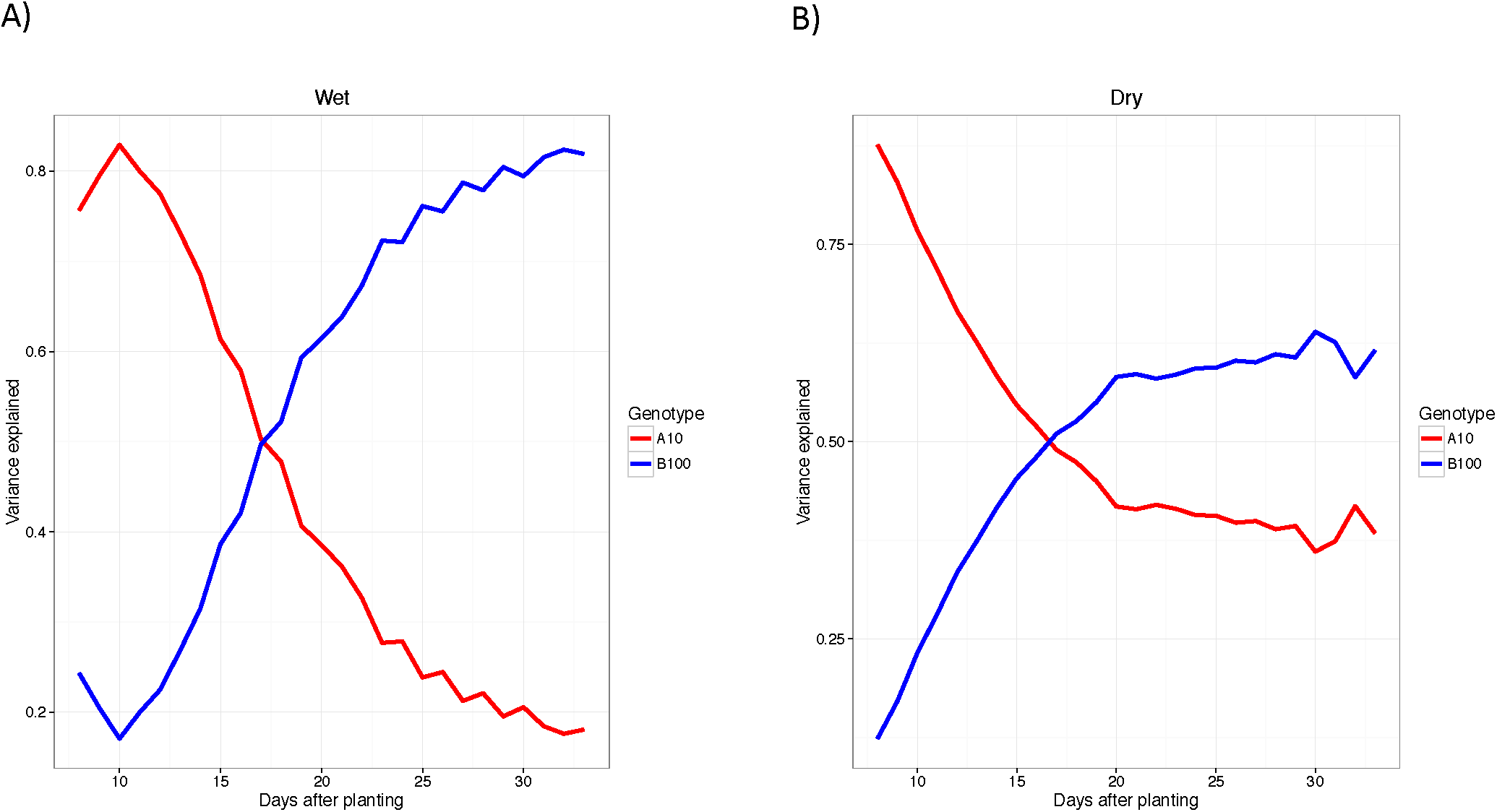
The proportional contribution of parental alleles that increase plant height changes throughout development during the 2014 experiment at Bellweather. Alleles derived from the A10 parent increase plant height early in development whereas the alleles that increase plant height later in development are inherited from the B100 parent. A) The proportion of additive genetic variance contributed by parental alleles plotted throughout all time points in the well-watered (wet) treatment block of the 2014 Bellweather experiment. B) The proportion of additive genetic variance contributed by parental alleles plotted throughout all time points in the water-limited (dry) treatment block of the 2014 Bellweather experiment.

Experiments performed in controlled environmental settings potentially end before the effects of the B100 alleles at positions 2@91 and 9@35 can fully materialize or may reflect limitations in growth due to pot size constraints. The results of a glass house experiment designed to test these factors, suggests that the pot size used during in the Bellweather trial does not dramatically limit vertical growth given the duration of this experiment (experiment ends prior to 35 days after planting; Figure 11). Clearly, pot size is a significantly influential factor for larger plants, later in development (Figure S14).

**Fig 11.**
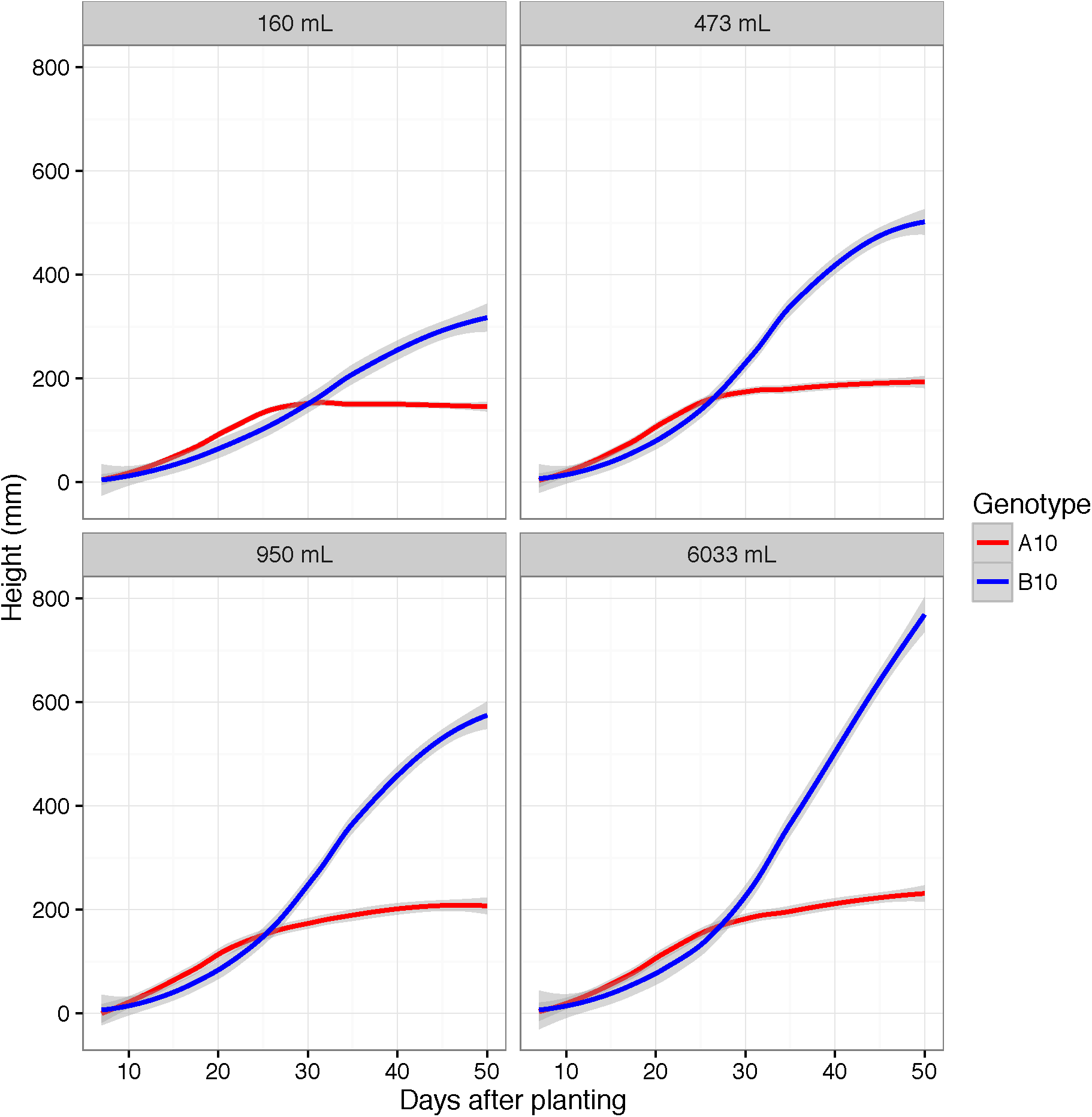
In replicated greenhouse experiments, the height of the A10 parent is greater than the B100 parent for the first 25 days a@er planting, a@er which the B100 parent becomes taller. Plants were grown in pots of four different sizes. Pots with volume greater than or equal 473 mL did not exhibit limita?ons in height at ?me points before 30 days aBer planting.

## Discussion

Although several end point analyses of plant height have been performed in grasses (7–14), we lack an understanding of the temporal interplay of genetics and environment over development. A major impediment is the challenge of performing replicated field trials that impose treatment effects and the difficulty of repeatedly measuring any trait manually. The use of model systems, robust field design strategies and high-throughput phenotyping platforms alleviate many of these obstacles.

The proportion of energy that grasses invest into vertical growth influences vegetative biomass and grain yield, which are important feed and biofuel traits (2,48,49). As demonstrated here, using platforms with high frequency and non-destructive measurements of plant height provides temporal resolution of QTL effects. The results of 12 coordinated grow outs of a *Setaria* RIL population in three different experimental locations suggest that plant height is specified as a functionally continuous developmental process that is under strong and robust genetic control but can be influenced by both water availability and planting density.

As expected (20–22), more aggressive water limitation treatments led to a greater reduction in plant height. However, no significant difference in the maximal plant growth rate was observed, rather it was delayed by approximately 3 days in water limited plants. Another surprising finding was that RILs planted at lower planting densities were taller than their densely planted counterpart, counter to the expectation that the shade avoidance response would result in increased plant height in densely grown stands (22,50) and may reflect resource limitations that inhibit growth of densely planted stands. Trial location was the most influential variable affecting plant height after genotype which may be the result of differences in day length and temperature influencing flowering time. In all field trials, flowering time was positively correlated with plant height at maturity.

Our results indicate that the identity of the genetic components that most strongly influence plant height are not dependent upon growth environment. Rather, delayed progression through this “core” genetic program that establishes mature plant height is a key element of G x E responses. Twelve genetic loci are found in each of the four different treatment regimens and are responsible for the strongest effects in every experiment, although many smaller effect QTL were identified across individual time points in multiple independent replicated trials. Overall, the identity of genetic loci identified in this study are highly congruent with an independent study of height performed on the same population (51).

The alleles that increase plant height are found in both parental lines, with the allele of largest positive effect (5@100) inherited from the shorter A10 parental line. The effects of this locus are realized early, as growth rate approaches its maximum value. Later in development, the number of loci that contribute to plant height increases. A vast majority of the loci that increase plant height late in development are inherited from the B100 parent. The dynamics of the late-appearing genetic loci are clearly dependent upon environmental factors as they are more variable between experiments.

Despite stark differences in rank order among genotypes and the relative height of the parents observed within field and controlled environments, the QTL detected largely overlap. The temporal QTL model provides the explanation for these conflicting observations across experiments conducted at different time windows. Plants grown at the Bellweather Phenotyping Facility are limited to the first 30-40 days of growth due to size constraints, whereas in the field data can be collected from more mature plants but cannot be reliably collected before 20 days after planting due to the need to establish seedlings after transplanting. These results demonstrate that adding temporal analysis to phenotyping can greatly improve our ability to detect and interpret the factors underlying complex traits.

The dense marker map created for this study can be combined with diverse germplasm studies to fine map causal variants and identify tightly linked markers for use in breeding programs. These tools can be used to modulate plant height and vertical growth rate throughout plant development given a predicted precipitation regimen or planting density. These approaches should be applicable across diverse species and traits for improvement of both food and biofuel crops.

## Conclusions

By combining field and controlled environmental analysis with quantitative genetics we were able to dissect the genetic and environmental components of plant height in *Setaria.* The identity of genetic components that determine plant height within this RIL population exhibits a strong degree of genetic canalization, but the contribution of these loci is influenced by temporal and environmental factors. Temporal analysis revealed that alleles from the wild parent act early in development, while alleles from the domesticated parent act later. This approach can be used to dissect all of the traits that can be measured in a high throughput manner by emerging phenotyping systems to better understand plant adaptation.

## Acknowledgements

This work was funded by the Department of Energy (DE-SC0008769) to the Donald Danforth Plant Science Center with sub awards to the University of Illinois and the Carnegie Institution for Science. This project would not have been possible without the dedicated efforts of the entire *Setaria* team (Baxter, Leakey, Dinneny, Brutnell, Rhee, Cousins and Voytas labs) and field crews (UIUC undergraduates and local high school students) who planted, nurtured and harvested over 100,000 plants over two years. Particularly, Mark Holmes, Hannah Schlake, Amanda Youssef, Sarah Keeley, Kara Barto, Jonny Yockey, Finey Ruan, Zack Reynado, Mitch Dickey, George Gunter, Marshall Alston-Yeagle, Audrey Rouse, and Andrew Chancellor for their collection of in-field measurements. We’d also like to thank Bradley Dalsing, Christopher Montes, Alex Hathcock, Kannan Puthuval and Dyson McMillan Singer the SoyFACE site technicians, for helping to maintain the experimental infrastructure. The authors are extremely grateful for their hard work. The authors would additionally like to thank Melinda Darnell, Malia Gehan and Noah Fahlgren for their assistance conducting the experiment at the Bellwether Phenotyping Facility and helpful discussions regarding analysis of the data.

